# A Multigraph model of the 3D genome

**DOI:** 10.1101/2021.11.11.468281

**Authors:** Diána Makai, András Cseh, Adél Sepsi, Szabolcs Makai

## Abstract

Spatial organisation of the genome has a fundamental effect on its biological functions. Chromatin-chromatin interactions and 3D spatial structures are involved in transcriptional regulation and have a decisive role in DNA replication and repair. To understand how individual genes and their regulatory elements function within the larger genomic context, and how the genome reacts as a whole to environmental stimuli, the linear sequence information needs to be interpreted in 3-dimensional space. While recent advances in chromatin conformation capture technologies including Hi-C, considerably advanced our understanding of the genomes, defining the DNA, as it is organized in the cell nucleus is still a challenging task. 3D genome modelling needs to reflect the DNA as a flexible polymer, which can wind up to the fraction of its total length and greatly unwind and stretch to implement a multitude of functions. Here we propose a novel approach to model genomes as a multigraph based on Hi-C contact data. Multigraph-based 3D genome modelling of barley and rice revealed the well-known Rabl and Rosetta chromatin organizations, respectively, as well as other higher order structures. Our results shows that the well-established toolset of Graph theory is highly valuable in modelling large genomes in 3D.

## INTRODUCTION

DNA in the living cell is organised into linear chromosomes that undergo dynamic architectural changes during the cell cycle, progressing from elongated interphase chromatin threads to condensed metaphase chromosomes ^1^. Within unreplicated chromosomes, a single continuous DNA molecule is tightly packaged into higher order structures. Nucleosomes form the chromatin fibre which are proposed to fold into consecutive chromatin loops ^2–4^ that are assembled into distinct chromosomal domains (topologically associating domains (TADs) in animals ^5,6^ and TAD-like domain structures in plants ^7,8^. The importance of chromatin packaging lays in its impact on DNA accessibility that consequently affects the fundamental processes of transcription, replication, and recombination. Changes in genome conformation can result in open chromatin regions serving as transcriptional hotspots while closed genomic regions have been demonstrated to be generally transcriptionally inactive ^9–13^.

The emergence of the high-throughput chromatin conformation capture sequencing (Hi-C) technologies has greatly contributed to the elucidation of spatial genome organisation in many organisms ^8,14,15^. However, determining the structure of the chromatin as it is organised in the cell nucleus is still a challenging task due to its extraordinary complexity and high plasticity. A Hi-C experiment sequences genomic fragments that are spatially co-located. By mapping the paired reads of chromatin capture sequencing to the reference genome, a contact matrix can be computed. Solving the contact matrix yields in the putative 3D structure of the genome ^16^. This approach represents a powerful tool to map simple, haploid genomes in 3D but appears more challenging when analysing complex, polyploid genomes. Chromatin capture sequencing cannot differentiate between the two parental genomes but chromosome representation can be interpreted as the average of the two homologous chromosomes ^17^.

Hi-C data allows us to make a model of the 3D genome up to a resolution of several thousands of base pairs ^18^. In these models, the genome is represented by bins of genomic fragments ranging in length between a few thousand and one million nucleotides. Kremer’s polymer modelling algorithm ^19^ has been frequently used to model DNA as a beads and springs. The method’s advantage is that it makes the model self-avoiding and computationally feasible. Kremer’s algorithm was originally developed to model polymers where every bead represents a monomer, and the springs represent the bonds between the monomers. Self-avoidance is achieved by setting the diameter of the individual spheres to be greater than the maximum length of a stretched spring. This constraint is reasonable for modelling polymers at a monomer level but may limit the biological relevance of Hi-C inferred 3D-genome models that are based on bins representing long DNA sequences.

The DNA is a flexible linear polymer ^20,21^, which in the living cell nucleus can wind up (like a bath towel) to the fraction of its total length and can unwind and stretch at a scale where transcription, double strand breaks and recombination occur ^22,23^. Genomic bins represent thousands of monomers and the length of the links between these bins exceed the diameter of the DNA fibre. For that reason, representing bins as monomers connected by short links limits interpretation of the 3D structure. Here we developed a novel approach to model and analyse large plant genomes in 3D, based on the graph theory of discrete mathematics. We portray the genome as a multigraph where Hi-C contact data is mapped to the chromosomes that are built up from sequentially adjacent bins identified from genomic data. This model allows the length of the bins while still using forces of springs that drives the 3D structure. Instead of point like beads, the bins represented by ‘elastic’ cylinders that add an extra dimension to the simulation notably that of the lengths of the individual bins (Figure 1). Self-avoidance in our modelling is implemented by dynamic checking for intersection/crossing of cylinders rather than artificially limiting the length of the links. These elastic cylinders behave like springs and can be stretched following Hooke’s law ^24^. Our technique is highly effective in modelling and visualizing the putative mechanical stress along the genomic region represented by the bin, where an elongated cylinder is interpreted as decondensed DNA, and a shortened cylinder represents a condensed DNA segment. In addition, by setting a threshold value to maximize the stretch forces, this technique allows to study the correlation between local genome conformation and the location of double strand breaks (DSB) along a chromosome. These parameters allow the control of the fibre stiffness during simulation.

**Figure 1:**
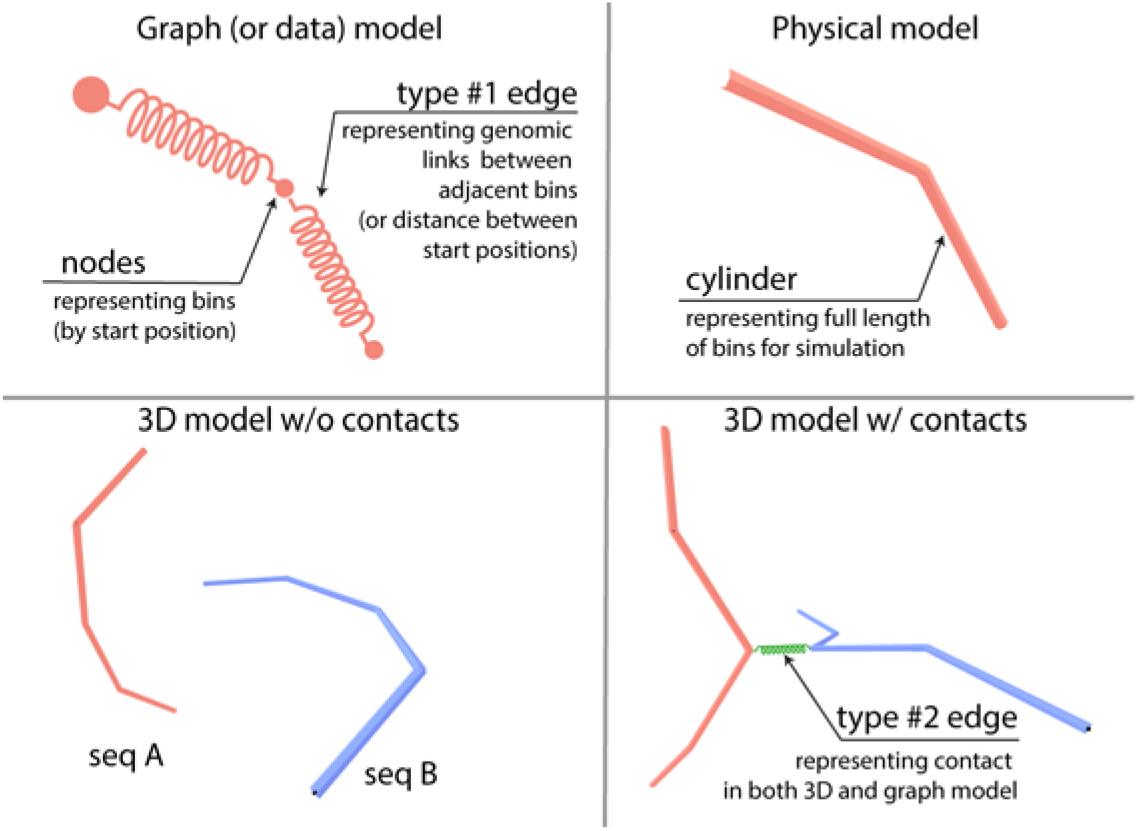
A visual summary of the multigraph model. The graph/data model depicts the genome as a graph of adjacent bins. The physical interpretation of the graph-based genome model is that nodes represent the start positions of the bins and edges are the physical representation of the bins that are visualized as cylinders. Two dummy chromosomes (seq A and seq B) without contacts set in an energy-minimalizing configuration. By adjusting the properties of the bins, the cylinders can be shorter or longer which in turn can be used as a model for condensed or decondensed state. Between nodes, a type #2 edge can be introduced based on contact data from Hi-C experiments. This puts extra strains on bins, therefore changing the configuration and length of cylinders (while its parameters are unchanged); hence, a graph based 3D genome. Since the genome and the contact graphs are two different graphs of the same node set (called vertices in graph theory), our 3D genome model is a multigraph.

In the present work our model explored bulk- and single-cell Hi-C data of two major cereal crops, barley (*Hordeum vulgare* cv. Golden Promise, 2n=2×=14, HH genome) and rice (*Oryza sativa* ssp *Japonica, 2n=2×=24*), to construct their 3D genome and carry out a comparative analysis between their topologies. We complemented our Hi-C modelling with three-dimensional cytological examination of barley chromosomes in cross-linked mitotic interphase nuclei, aided by optical sectioning and high-resolution confocal microscopy.

## MATERIALS AND METHODS

### Raw data analyses

The Hi-C library of barley (NCBI Accession No. SRR8922888) used in this study is from one week old seedlings of cv. Golden Promise ^25^. The single-cell libraries of rice (NCBI Accession No. SRR8261296 and SRR8261290, respectively) used here are from shoot cell isolated from seedlings 12-day-after-germination and from sperm isolated from rice stamens ^26^. Hi-C data was processed following the method described by Padmarasu and co-workers ^27^. We used high-quality ‘Golden Promise’ reference assembly ^25^ and ‘*Japonica*’ reference IRGSP-1.0.

### Contact statistics and pre-filtering

Population averaged (multi-cell) barley library resulted in over 20 million interactions. We used a 10 kbase resolution that reduced the interchromosomal contacts to 6.5M and the intrachromosomal contacts to 5M. A contact distribution between the 10 kb bins was calculated and a percentile-based threshold of 85 for contact counts was determined to select strong interactions (Figure 2). This number corresponds to percentile of 0.512 and 0.524 order for intra and interchromosomal respectively.

**Figure 2:**
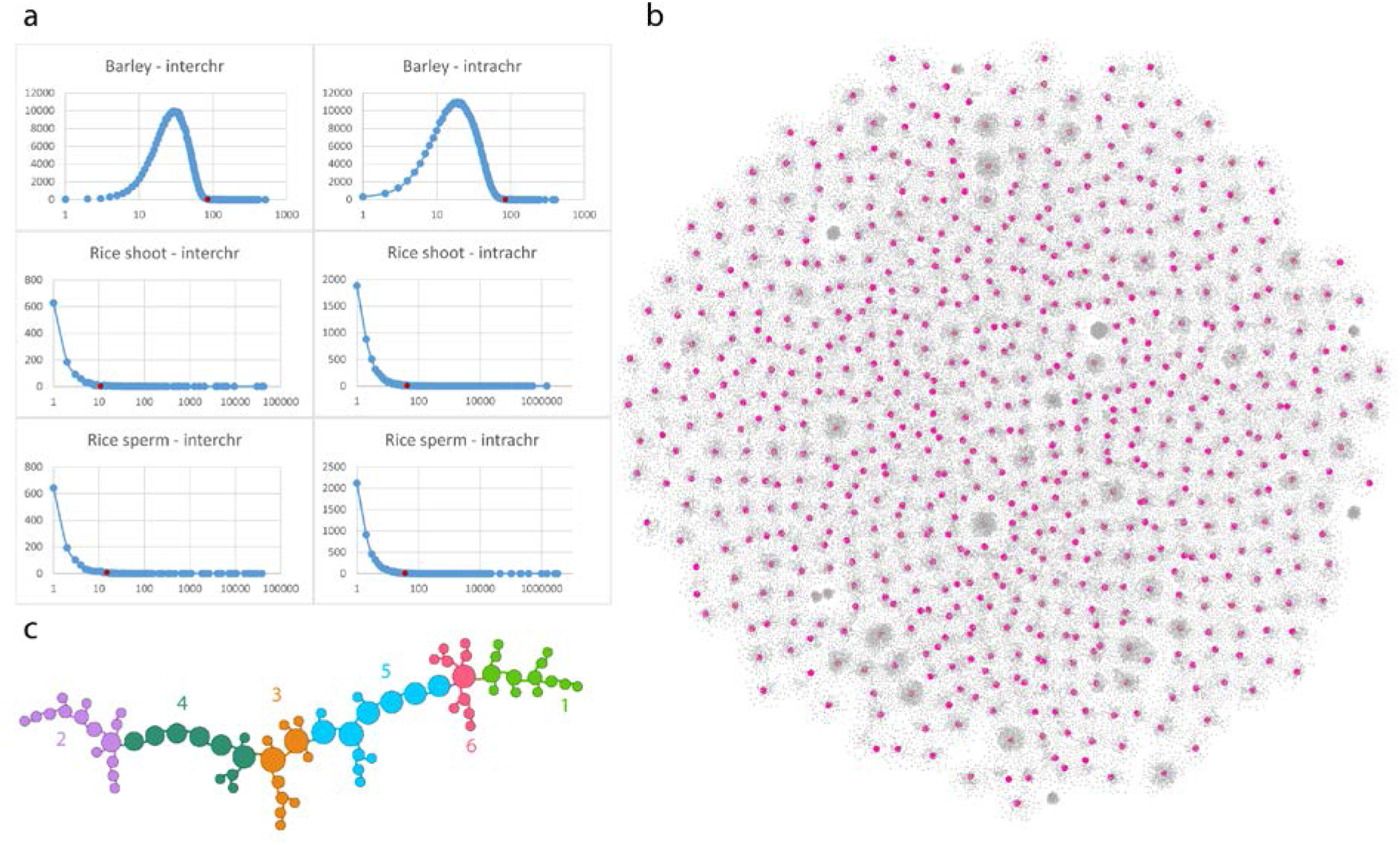
Contact graph construction and analysis. (a) Contacts per bin distribution of selected Hi-C libraries. The curve shows the number of bins that has a given contact count. The x-axis is logarithmic, and shows contact count, the y-axis is linear and shows the number of bins. On the charts, a red dot shows the cut-off value for each analysis (see numbers in the M&M section of the manuscript). For further downstream analysis, only bins that have a frequency of this value or higher were used. In all cases self-loops were discarded and not counted. (b) 2D representation of interchromosomal filtered contacts of Barley (See methods). Magenta nodes represent bins that have 14, or more than 14 edges in the graph. The layout indicates that contacts between the non-homologuous chromosomes are organized by dedicated genomic regions, suggesting hierarchical genomic structures. (c) The largest subgraph after filtering for bins of 14 or more different contacts. An intriguing linear layout of contacts has emerged suggesting a signal transduction „highway” that might function like Newton’s cradle. Colors represent the clusters of the graph. The clusters were characterized by gene enrichment analysis that revealed housekeeping genes within the regions represented by the bins.

The two single cell libraries of rice presented fewer contacts. The shoot cell library had 105,649 interchromosomal and 5.6M intrachromosomal contacts. Similarly, the sperm cell library had 102,780 interchromosomal and 10.3M intrachromosomal contacts at 10 kilobase bin size. The data was further filtered based on percentile data. Cutoff values of 11 and 43 for shoot cell and 11 and 36 for sperm cell were used for inter- and intrachromosomal edges respectively (Figure 2a). Intra-bin contacts (self-loops) were discarded in all cases in our analysis.

### Multigraph construction

Our model is based on the classic beads on string polymer model of Kremer et co-workers ^19^. This model was originally proposed to model polymers on monomer level. Self-avoidance was achieved by not allowing links between beads longer then the radius of the beads. This implies that bins are not flexible and supposes to be in a uniform state, which presumes DNA as a rigid molecule with a static structure. *A modification, therefore, is necessary because a single bead in our model is a bin that represent ten thousand nucleotides (monomers)*.

In our graph structure, nodes represent the starting position of the bins. Between nodes representing sequentially adjacent bin positions, an edge represent the bin itself. We refer to this category of edges (type #1 edge on Figure 1) as genomic edges, since they represent genomic regions in a sequential order that ultimately form an entire chromosome. These edges can be represented as elastic cylinders whose length may indicate the strain present in the DNA region. According to the graph structure, a chromosome without contacts can be visualised as a single, untangled rope. Subsequently, Hi-C derived contact data is mapped to the graph that adds the edges between nodes of interacting bins. Since contacts are studied at bin resolution, drawing contacts between the ‘start position’ is sufficient. We refer this type of edges as contact edges (type #2 edge on Figure 1). Hence, the 3D genome is displayed as a multigraph considering that the nodes can be linked by two different types of edges: genomic edges and contact edges (Figure 1).

The calculation of the 3D structure was based on a physical model of springs ^28^. The forces considered were: i) repulsion between cylinders, ii) repulsion between the two ends of the cylinders (driving decondensation), iii) spring force in cylinders (driving condensation) and iv) drag force of the environment. The algorithm was run to minimalize “energy” of the springs/cylinders. In addition, the length of cylinders were adjustable to model condensed and decondensed state of the chromatin; a shorter cylinder thus packs the same amount of base pairs in shorter section of the strand. The diameter of the cylinders was constant, as we did not intend to model this aspect of the chromatin. An additional constraint on node positions was applied in a shape of a sphere that represents the nucleus.

Pure contact graphs were modelled by graph analysis as described by Das and co-workers ^17^. The 2D modelling of graphs were performed by Gephi ^29^. The 3D multigraphs were generated by our own tools using webGL, NodeJS technology and ngraphs ^30^.

### Cytological analysis

*In situ* hybridization was carried out on mitotic root tips nuclei of barley cv. Golden Promise according to the protocol described by ^31^ by using the barley centromere-specific G+C sequences ^32^ and the plant universal telomeric repeat sequences (TRS) as probes ^33^. Probe labelling was carried out by Nick-translation (AF594 NT Labeling Kit, PP-305L-AF594; AF488 NT Labeling Kit, PP-305L-AF488; respectively; Jena Bioscience, Jena, Germany). Slides were counterstained in 12 μl of Vectashield Antifade Mounting Medium with DAPI (H-1200; Vector Laboratories, Burlingame, CA, USA).

ImmunoGISH/FISH was performed on mitotic nuclei of a wheat-barley 7BS.7HL disomic translocation line ^34^ obtained from tapetum cells of the anther tissue. Paraformaldehyde crosslinked mitotic nuclei were slide mounted and the ImmunoGISH/FISH procedure was carried out as described previously by Sepsi et al. (2018). To localise active centromeres, immunolabelling was carried out with a rabbit anti-grass CENH3 antibody ^35^ detected by a goat anti-rabbit abberior Star Red (STRED-1002-500UG, Abberior Instruments, Göttingen, Germany) secondary antibody. Simultaneous *in situ* hybridization included Nick-translation labelled genomic DNA from barley (AF594 NT Labeling Kit, PP-305L-AF594, Jena Bioscience) and the TRS-probe (AF488 NT Labeling Kit, PP-305L-AF488).

Confocal microscopy was performed using a TCS SP8 confocal laser-scanning microscope (Leica Microsystems GmbH, Wetzlar, Germany). Z stacks were acquired using a HC PL APO CS2 639/1.40 oil immersion objective (Leica Microsystems GmbH). Image acquisition was carried out by bidirectional scanning along the x-axis, and images were averaged from three distinct image frames to reduce image noise. The fluorochromes used in the present study were: DAPI (excited at 405 nm, detected from 410 to 450 nm), Alexa Fluor 488 (excited at 488 nm, detected from 490 to 520 nm), Alexa Fluor 594 (excited at 561 nm, detected from 565 to 620 nm).

## RESULTS

For both single cell and bulk (population averaged) processed Hi-C data a contact graph was built by using bins as nodes and contacts as edges. We added all bins of each chromosome to the graph, even those that were not participating in any contacts. These nodes only had one or two edges linking them to nodes representing neighboring bins on the chromosome. The resulting graph was a multigraph that contained a graph representation of the full genome integrated with the contact graph. In other words, two types of edges can link a node in the graph, (i) genomic link between nodes representing (sequentially) adjacent bins and (ii) contacts of Hi-C detected interaction that can be either intrachromosomal or interchromosomal (Figure 1). The adjusted graph layout algorithm calculates the 3D layout of the multigraphs following the physical model of springs and we propose that the resulting structure can be interpreted as an improved 3D genome model. Contact frequency was additionally used to tune the elasticity value of the springs representing the contacts. Higher frequency rate rendered the spring more rigid, implying that it stretched less and thus resulted in shorter segments.

### Barley (multicellular, bulk samples)

Population samples can provide rich data that are, on the other hand, more difficult to interpret ^36^. We used graph analysis tools to obtain a better understanding of the contact data gained from mixed cell types of 10 days old seedlings of barley. First, we present the pure contact graphs of the barley Hi-C data, and we subsequently introduce the multigraph-based 3D genome model.

#### Pure contact graphs of barley

To reduce complexity, we built contact graphs for interchromosomal and intrachromosomal contacts separately. Both cases demonstrated very complex graphs with non-random layouts (Figure 2b, 2c, 3a).

**Figure 3:**
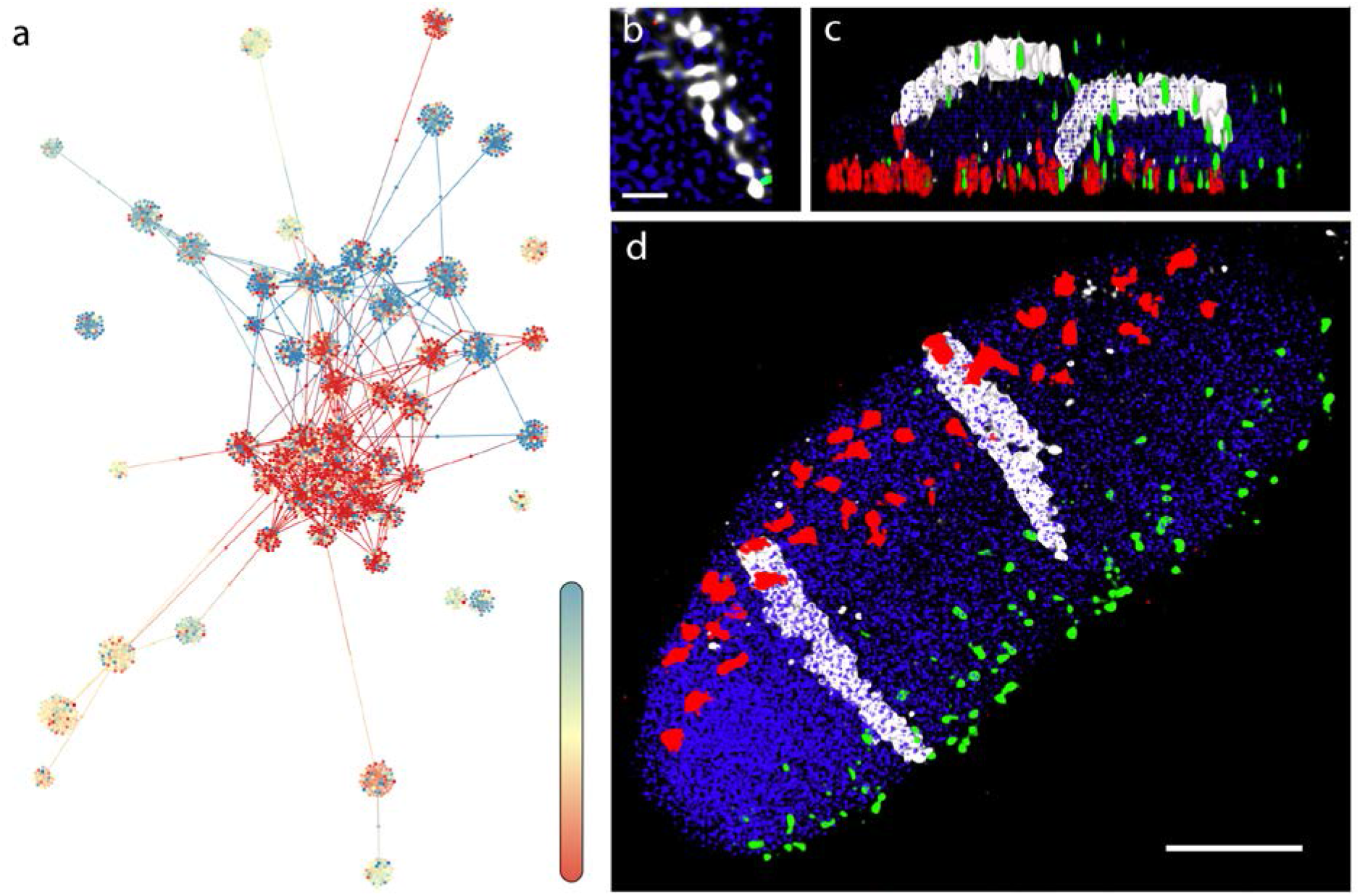
2D graph representation of contact data of chromosome 7H of barley and Microscopic evaluation of the organization of an individual (long) arm of the 7H barley chromosome (7HL) within the interphase nucleus. a) 2D graph representation of contact data of chromosome 7H. Contacts are filtered for intrachromosomal and 85 hits or above. Fully saturated blue and red colours represent the two telomeric ends of the chromosome. Positions are indicated by a gradient colour in between the ends, where pale yellow marks the putative location of the pericentromeric region. Contacts are clustered within each telomeric regions with a few contacts between them. The pericentomeric region is well separated from the telomeres, a contact pattern typical of the Rabl configuration. b) A high-resolution image enlargement of the 7HL chromatin threads (white) intertwined with the neighboring chromatin captured within a mitotic nucleus of the 7BS-7HL wheat-barley translocation line with confocal laser-scanning microscopy. The 7BS-7HL wheat-barley translocation line carries the 7HL barley chromosome arms stably transferred into the wheat background, allowing the 3D visualization of the barley chromatin by genomic *in situ* hybridization (GISH). c) Side view of a three-dimensionally rendered z-stack captured from a mitotic nucleus of the 7BS-7HL wheat-barley translocation line. The 7HL barley chromatin is detected by GISH (shown in white), centromeres are visualised by simultaneous immunolabelling (red) and telomeres are shown by fluorescence in situ hybridisation (green). d) Front view of the same three-dimensionally rendered z-stack, showing parallel arrangement of the two homologous 7H long arms. Rabl organization is clear from the polarized centromere (red signals)-telomere (green signals) arrangement and the signal of the perpendicularly running barley chromosome arms. Bar=5um, except Figure 3b where bar represents 1⍰⍰m

The **interchromosomal** contact graphs showed 73 738 bins and 82 807 contacts (Figure 2b). Using the 2D OpenOrd layout algorithm ^37^, the graph demonstrated a clustered layout with a few bins that were involved in a high number of contacts while the majority of bins had only contacts to a single bin. This prompted us to filter nodes by their degree. A cutoff value of 14 for degree was set. The result presented 371 bins, 297 edges, and a topology of many unlinked subgraphs. Subsequently, we analyzed the greatest subgraph that demonstrated a chain-like layout of bins (Figure 2c). Given that the first filtering (percentile based) filters for bins participating in **over 85 contacts** and second filtering further filters these bins for those that participate in at least 14 *different* contacts, we suggest that the subgraphs represent highly conservative contact paths. Gene enrichment analysis of encoded genes within these bins and in their proximity showed an over-representation of housekeeping genes (Supplementary Table 1). Since the experiment was carried out on bulk samples of 10 days old seedlings, we propose that this network of interacting bins is conserved across all cell types.

When applying the 2D OpenOrd algorithm to bins of a single chromosome with over 85 **intrachromosomal** contacts, the emerged modular layout displayed topologically distinct node clusters, indicative of TAD-like structures (Figure 3a) ^8^. For instance, a 2D graph for the 7H chromosome (Figure 3d) shows nodes at centers of flower like conformations further demonstrating that these are TAD like structures. The colouring of the nodes indicates the position of the bin within the chromosomes. The two terminal segments (telomeres, blue and red) are well separated and only a few edges are drawn across the blue and red domains, which shows a low number of intrachromosomal contacts between the distal regions detected by the Hi-C analysis. Moreover, on this topology a third region, located proximally, far from the telomeres (i.e. pericentromeres) is well separated from the rest of the chromosome (Figure 3a). This characteristic layout, where the distal and proximal regions showed a polarised arrangement appears to depict the highly conserved Rabl chromosome configuration ^38^ characteristic of barley interphase chromosomes ^39^.

#### Cytological visualisation of the spatial organisation of a single 7H barley chromosome arm

A mitotic nuclei of a wheat/barley introgression line was used to show the organization of a single barley chromosome arm in the interphase nucleus. The barley chromosome arms have been detected by GISH while telomeres were visualized by FISH using the TRS-probe. Active centromeres were shown by simultaneous immunolabelling for the grass-specific centromere-specific Histone H3 protein (CENH3). The fine structure of the 7H barley chromatin showed chromatin threads as a scaffold, interdigitated with the neighbouring chromatin fibres (Figure 3b). Optical sectioning revealed clear polarization of the telomeres and centromeres in opposite nuclear hemispheres while the barley 7H chromosome arm pair run perpendicular between the two nuclear poles (N of nuclei=55) and occupied separate territories (Figure 3b-d).

#### Multigraph representation of the 3D genome

To build the multigraph model of the 3D genome, we first merged all intrachromosomal (Figure 4a, left) and the interchromosomal contact graphs into a single, combined contact graph (Figure 4a, right). We then integrated the combined graph with the genomic graph where the bins represented by nodes were arranged sequentially to ultimately form the individual chromosomes, so the added edges represented genomic links (type #1 edge, Figure 1) between the adjacent bins. These newly added nodes determined the positions, and their outgoing edges determined the orientations, lengths, and stresses of the cylinders. Each edge of a genomic link is checked against intercrossing with other, non-adjacent edges. In the multigraph, the contact edges (type #2 edge, Figure 1) are not drawn, they can intercross, and they do not snap.

**Figure 4:**
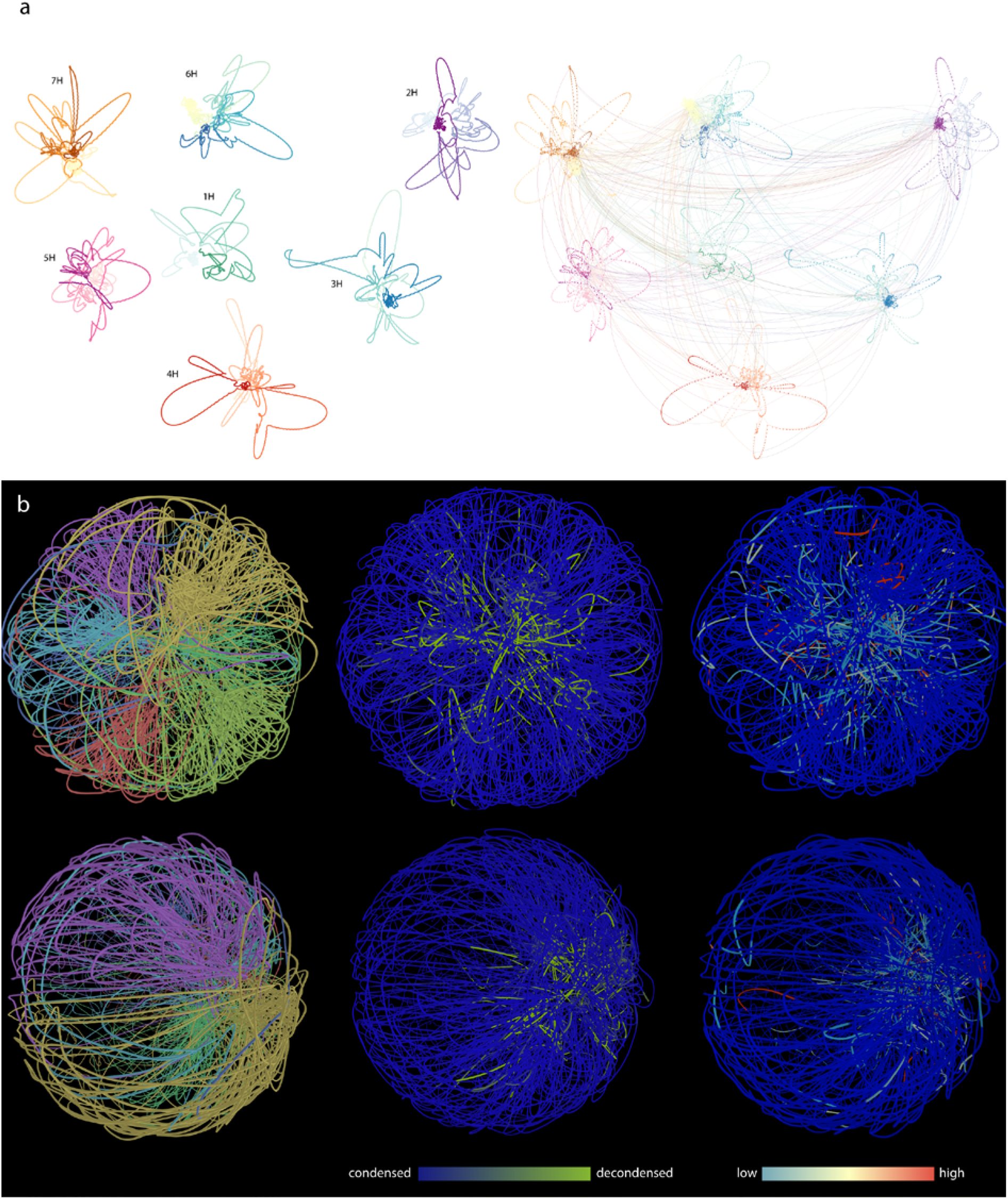
Going from 2D contact graph to multigraph model of 3D genome of barley. (a) 2D layouts of the multigraph of each barley chromosomes. Layout was calculated based on the intrachromosomal contacts only. Entanglement of the two ends of the chromosomes are clearly visible showing the Rabl configuration. Right panel: interchromosomal contacts are added. (b) Barley 3D genomic model. The two rows represent two orthogonal perspective of the genome model. Colouring is based on different attributes. On the left panels, chromosomes are colour coded. On the central images, condensation/decondensation is visualized by blue and green colours, respectively. On the right panels, expression data of root tissue (source: the PRJEB14349 bioproject) is mapped to the 3D genome model.

In the 3D layout, well-defined chromosome territories were observed where the chromosomes clearly folded in half (Figure 4b). Every simulation was inherent to the initial state, therefore the final layout changed as we modified the initial state of the multigraph. However, independent of the initial state, chromosomes always folded in half so that the telomeric region of each chromosomes clustered together, making a nucleolus-like, cavernous (hollow) formation spontaneously emerging during the simulation (Figure 5d). This conformation was off center, pushed towards the perimeter of the nucleus depending on the initial changes and/or the modifications of different parameters during the simulation. In cases when this formation was smeared along the nucleus, an architecture reminiscent of the experimentally observed Rabl configuration appeared (Figure 5).

**Figure 5:**
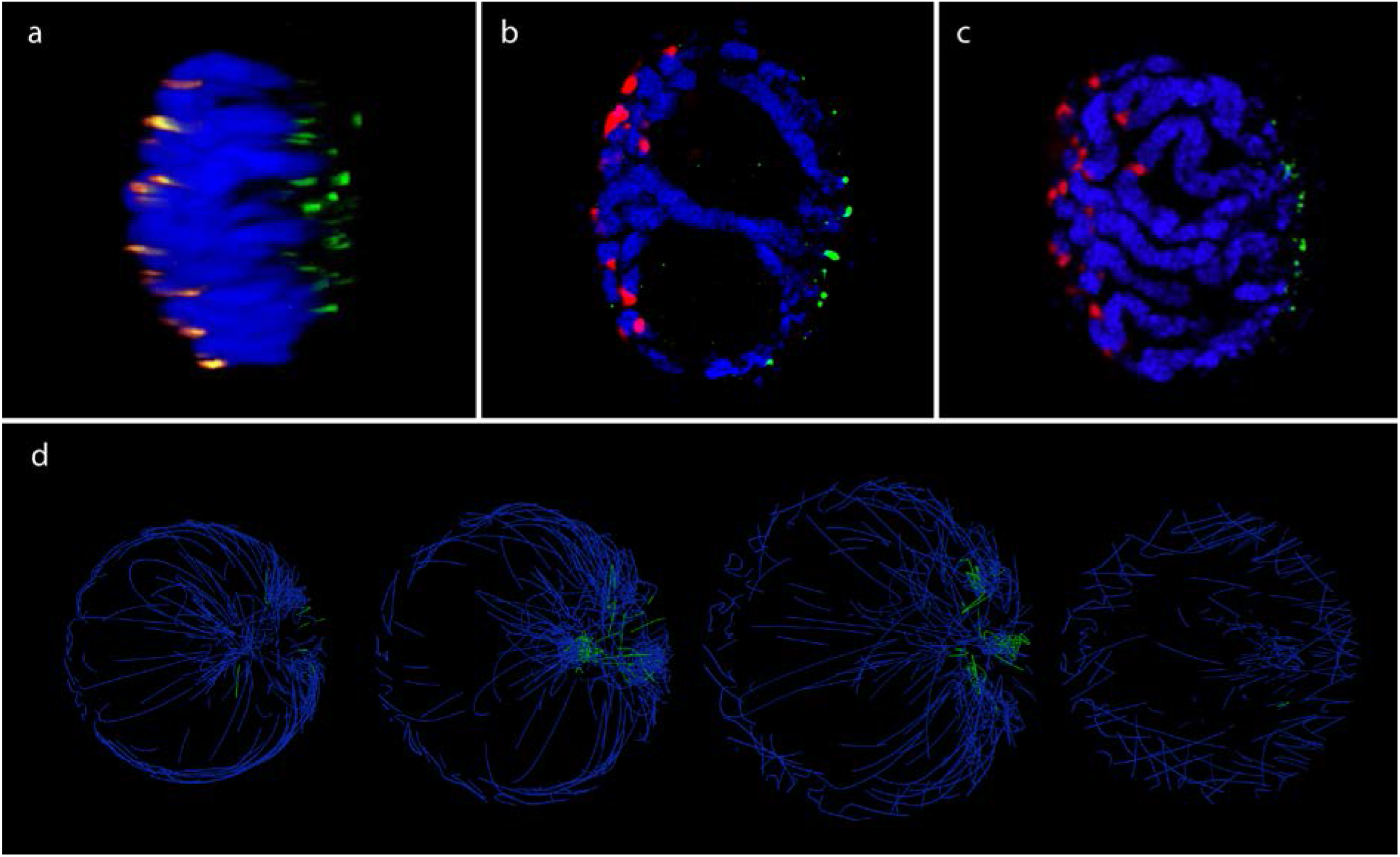
*In situ* and *in silico.* Fluorescence *in situ* hybridization (FISH) showing centromeres (orange, red) and telomeres (green) on a 3D-fixed barley mitotic nuclei. Barley centromeres are labelled by the barley centromere-specific G+C repeat probe (red) while telomeres are visualized by the plant telomeric repeat sequences (TRS) used as a probe (green). Chromosomes are counterstained with DAPI. Bar=5 μm. a) Side-view of a 3D-rendered z-stack captured from a barley mitotic nuclei. b)-c) FISH on individual image frames of a barley mitotic nucleus showing polarized centromere-telomere arrangement and chromosome arms laying parallel with the telomere-centromere axis. d) *In silico*, cross section images of the Barley 3D genome model. Green indicates telomeric regions. A well-distinguished nucleolus-like region is visible inside the spherical topology. Telomeres are clustered on one side of the spherical model.

The resulting layout not only gave a 3D model of the genome, highly similar to that observed under the microscope (Figure 5b), but also showed the map of the mechanical stress along the chromosomes (Figure 4b). Mapping expression data along the bins correlated with the elongated state of the cylinders, as we found most of the transcriptionally active spots on elongated cylinders (Figure 4b).

#### 3D visualization of chromosome arrangement in barley cv. Golden Promise mitotic interphase nuclei

Our microscopic 3D imaging of barley cv. Golden Promise mitotic interphase nuclei revealed similar nuclear organization of chromosomes (Figure 5a-c). Barley telomeres were shown by FISH with the TRS probe while barley centromeres were visualized by the barley centromere-specific G+C repeat sequences. The chromatin was contrast stained with DAPI (Figure 5a-c). According to these experiments, centromeres and telomeres were polarized in the opposite hemisphere of the nucleus in all cases (N=45, Figure 5a-c) while the chromatin run perpendicular as parallel threads, enclosing one/two centrally located large nucleolus (Figure 5b).

### Rice (single cell experiments)

We applied our 3D genome modelling technique on diploid and haploid single cell data of rice. We used *sc*Hi-C datasets of rice generated by Zhou and co-workers ^26^.

We selected a haploid sperm cell and the diploid shoot cell libraries to implement our model. The first was chosen as a ‘vanilla’ experiment with ideal dataset: haploid and single cell. The second was chosen to serve as a basis of comparison to the barley experiments carried out on similar cell type. Yet, a large variance of count number is displayed that we assume to be the effect of biases of amplification and sequencing ^40^.

#### Pure contact graphs of rice

The interchromomal contact graph of the shoot cell resulted in 802 nodes and 1501 edges while the intrachromomal contact graph had 347 nodes and 255 edges after the percentile based filtering. The merged contact graph had 1009 nodes and 1756 edges. The contact frequency showed a huge variance. The interchromomal contact graph of the sperm cell had 1063 nodes and 1861 edges while the intrachromomal contact graph had 161 nodes and 122 edges after the percentile based filtering. The merged contact graph had 1125 nodes and 1983 edges. The contact frequency showed a huge variance. The interchromosomal graphs of the cell types had different topology (data not shown).

#### Multigraph representation of single cell Hi-C data of rice

The intrachromosomal contacts were rather sparse whereas interchromosomal contacts were better represented for both shoot (Figure 6a, 6b) and sperm cells. Simulation drew a graph of the chromosomes that was tightly entangled. Most of the contacts concentrated in the middle of the sphere, to the estimated location of the nucleus. The high concentration of interchromosomal contacts formed a cavity in the middle that corresponded to the approximate nuclear position of the nucleolus (Figure 6e, f).

**Figure 6:**
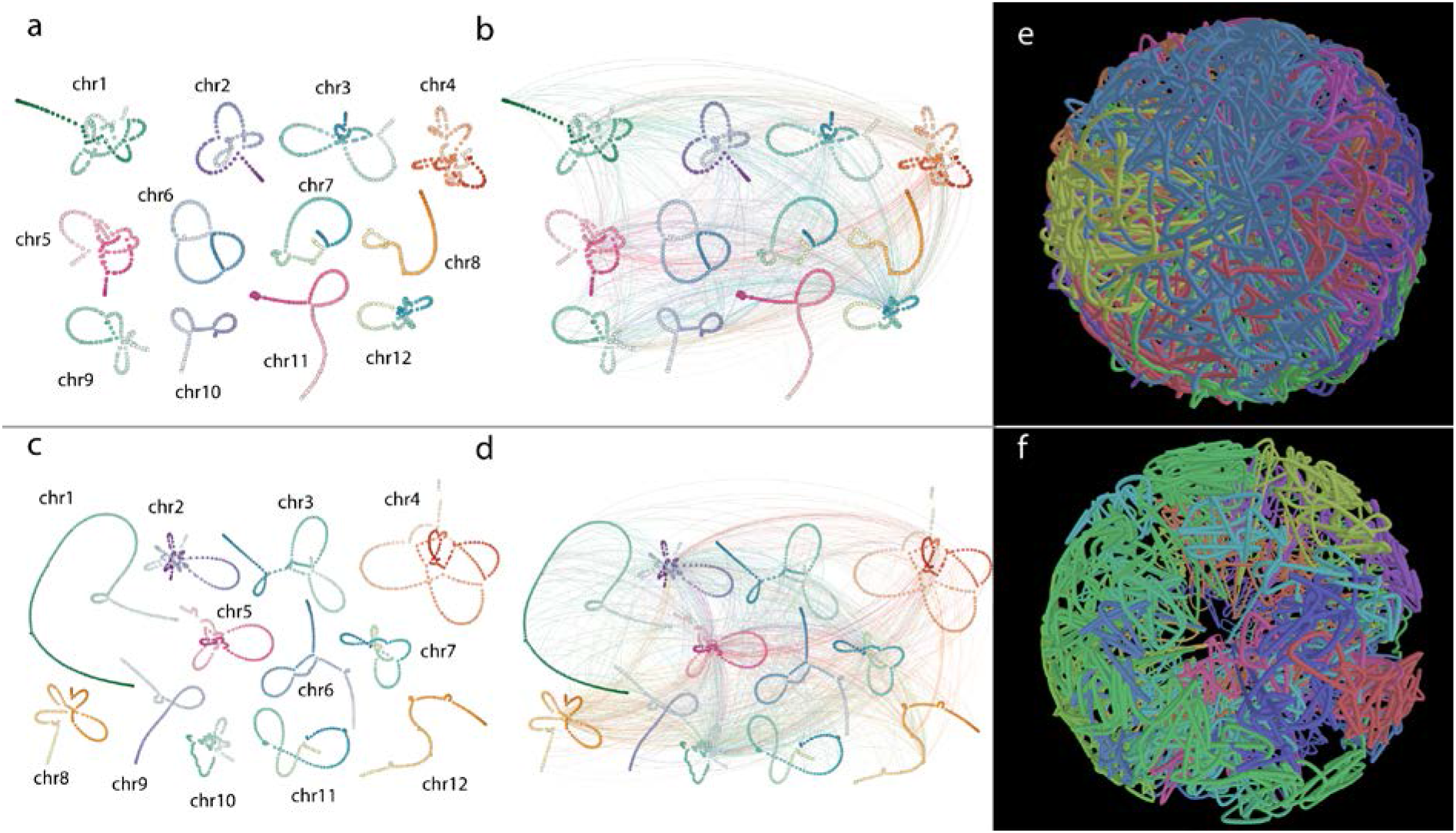
Rice *sc*Hi-C data-based genome graph construction. (a, b) The 2D multigraph layout of rice shoot cells. Left without interchromosomal contacts (a), right interchromosomal contacts drawn (b). (c, d) The 2D multigraph image of rice sperm cell. Left without interchromosomal contacts, right interchromosomal contacts drawn. (e) 3D genome of rice shoot cell based on single cell Hi-C data. (f) 3D genome model of rice sperm cell. Chromosome colour coding shows apparent chromosome territories. Cavity in the centre of the sphere occurs spontaneously during simulation, suggesting the putative location of the nucleolus.

#### Comparative analysis of barley and rice Hi-C data

The genomes of rice and barley differ in both size and their reported configuration. We therefore chose their available shoot cell data libraries to compare the 3D-genome models generated by our analysis. Both the 2D and 3D representations of the intrachromosomal multigraphs showed distinct features for barley and rice. Individual barley chromosomes were entangled by their two ends and had large loose loops in between (Figure 4 and 5). In the case of rice, chromosome ends appeared to be less interactive, however more loops formed around a center that was reminiscent of a flower structure (see supplementary videos). Expected, but equally surprising that both cases corresponded to the of the historically reported Rabl- and Rosetta conformations of barley and rice. Graph tools outlined these topologies without applying any opinionated and/or arbiter filtering besides filtering to reduce noise.

## DISCUSSION

Genomes can be perceived as graphs, a network of genomic bins described by a binary adjacency matrix. In this matrix only adjacent bins have links; all other elements of the matrix are 0. Hi-C analysis also results in an adjacency matrix that is based on captured genomic interactions. In this matrix, the values represent interaction frequencies between the bins (0 means no interaction, any positive integer shows the count of measured interaction). The two graphs (contact graph and genomic graph) can be merged into a multigraph and an approximation of the genomic configuration can be modeled by means of graph theory tools.

The chromatin 3D structure in the nucleus is assumed to be determined by its function and not by its thermodynamic equilibrium ^41,42^. Elements of higher-order chromatin organization and chromatin dynamics show a considerable flexibility during the cell cycle. Specific features of chromatin architecture may vary from species to species and cell to cell while conserved phenomena also exist. For instance, in plant nuclei, interphase chromosomes are organized in distinct territories ^43,44^ where landmark chromosomal regions have a well-defined orientation. A highly conserved aspect of chromatin organisation in large genome cereals, such as wheat and barley, is the orientation of their telomeres and centromeres into opposite poles of the nuclear periphery in a so-called “Rabl” configuration ^45,46^. Rice nuclei in contrast, carrying the smallest genome among the grasses ^47^, show “Rosette” configuration where pericentromeric repeat elements are located in the proximity of the nucleolus from which coding regions emerge as chromatin loops ^46,48,49^. Both type of configuration has its characteristic contact pattern.

In this paper, we demonstrated the efficiency of Graph theory-based tools in constructing 3D genomes from Hi-C libraries of barley and rice. One advantage of the multigraph approach is that it allows the combination of two independent datasets (chromosome sequence data and Hi-C data) into one single data model. During the layout calculation of the graph, the binary adjacency matrix puts a constraint on bins to keep their sequential order intact although at varying length. The interaction-based adjacency matrix constrains the bins by the “captured” conformation. Both constrains influences the final confirmation of the genomes, however only the genomic graph is drawn to visualize the 3D genome.

Another advantage of the presented model is that it allows the stretching and shrinking of the bins, thus simulating strains along the DNA. Yet, the greatest advantage is that representing the whole genome as a graph brings the well-established and proven toolset of Graph theory on board of the Hi-C based 3D genome modeling. Mathematical models, layout algorithms, similarity measures, clustering are just a few of the tools that can be reused from Graph theory in Hi-C data analysis and 3D genome reconstruction.

The contact graph analysis of rice indicated a conformation similar to that of earlier reported as KNOT conformation by Grob and co-workers ^8^. In barley, filtering of the contact graph highlighted a single contact pathway that resembles to Newton’s cradle (Figure 2). Such linear contact path may hint evidences to the mechanistic principles governing the genome ^50,51^. We examined the genes that are typically abundant within these locations and found overrepresentation of house-keeping genes. While this result is in line with expectations, GO enrichment analysis tends to highlight HK genes as they are the most accurately (thus most frequently) annotated genes. Nevertheless, it is reasonable to think that a subset of bins that form a conserved interaction network across cell types carry genes with conserved functions. This further elucidates that these bins may be constitutively active transcriptionally and bear a decondensed, open chromatin structure.

In our 3D model most of the interchromosomal contacts gathered around a naturally occurring cavity, resembling a nucleolus in living cells. This might suggest that activity shaping the genome are present at nucleolus perimeter and that and Hi-C may be particularly sensitive to contacts around the nucleolus. However, more Hi-C analysis and comparison is needed to confirm this. The formaldehyde treatment used by the Hi-C technology to fix hydrogen bonds may equally capture temporarily collocated regions, such as enhancer-promoter interactions and more stable connections, such as euchromatin regions localized in the proximity of the nuclear envelope or the nucleolus ^52^.

Another compelling insight was offered by the fact that our method allowed the observation of chromosome dynamics. We noted that certain chromosome conformations become locked up and cannot be untangled during metaphases, when chromosomes need to separate and migrate to opposite poles of the cell. While it is well probable that our model does not consider all parameters acting on the DNA molecules *in vivo*, and the available data is scarce, we allowed cylinders, representing the bins of the genome to break. This condition alone facilitated the untangling of all conformations tested by our experiments. Our results highlight the effect of DNA breaks on the topological complications caused by the processing of the DNA double helix during the cell cycle. The multigraph-based genome model accentuated the necessity of DNA cleavage in order to allow quick disentangling of chromosomes. These insights may hint that double strand breaks introduced into the DNA phosphodiester backbone and the subsequent repair mechanisms may have a role in facilitating rapid chromatin condensation. Thus, DSBs and repair mechanisms may have offered an evolutionary advantage by accelerating the cell cycle. Once sexual reproduction emerged, recombination may have arisen from this very same process, bringing further evolutionary advantages.

## DATA AVAILABILITY

The datasets generated and analyzed during the current study are available from the corresponding author upon reasonable request.

## ETHICAL STANDARDS

On behalf of all co-authors, the corresponding author states that the work described is original, previously unpublished research. All the authors listed have approved the manuscript.

## ACKNOWLEDGEMENTS

This research was funded by research grants from the National Research, Development and Innovation Office (NKFIH-FK-124266 and NKFIH-K-129221).

## AUTHOR CONTRIBUTIONS STATEMENT

SM, AS and AC conceived the designed the experiment. DM and SA conducted laboratory experiments. SM, AS, AC and DM carried out all data analyses and prepared the figures. SM and AS wrote the original manuscript and all other authors reviewed and approved the final manuscript. AS and AC provided resources and obtained funding for the work.

## SUPPLEMENTARY MATERIAL

Supplementary Table 1: The result of the GO enrichment study for the clusters shown on Figure 2c.

Supplementary Video 1: Animated 3D view of the multigraph 3D model of barely. Chromosomes are color coded.

Supplementary Video 2: Chromosome 7H of barley highlighted in the model to show the polarized, Rabl like conformation.

Supplementary Video 3: Chromosome 3 of rice shoot cell highlighted in the model to show the Rosetta like conformation.

Supplementary Video 4: Chromosome 8 of rice shoot cell highlighted in the model to show the Rosetta like conformation.

Supplementary Video 5: Chromosome 1 of rice sperm cell highlighted in the model to show the Rosetta like conformation.

Supplementary Video 6: Chromosome 2 of rice sperm cell highlighted in the model to show the Rosetta like conformation.

Supplementary Video 7: Chromosome 8 of rice sperm cell highlighted in the model to show the Rosetta like conformation.

